# Nutrient loading from a sustainably certified aquaculture operation dwarfs annual nutrient inputs from a large multi-use watershed, Lake Yojoa, Honduras

**DOI:** 10.1101/2024.03.02.583117

**Authors:** J.M. Fadum, M.V.R. Ross, E. A. Tenorio, C.A. Barby, E.K. Hall

**Author notes:** Corresponding authors: Jemma Fadum  and Ed Hall. Jemma Fadum is currently at Carnegie Institution for Science in the Department of Global Ecology, Stanford, CA.

## Abstract

Net-pen aquaculture is a popular and increasingly prevalent method for producing large quantities of low-fat protein in freshwater ecosystems across the tropics. While there are numerable social and economic advantages associated with aquaculture, there are also challenges related to the environmental sustainability of aquaculture. Chief among these risks is excessive nutrient loading which can drive rapid eutrophication in aquatic ecosystems. In this study, we compare the estimated annual nutrient loads from the six principal tributaries that contribute to Lake Yojoa to the estimated nutrient load of a large net-pen Tilapia operation located in the central region of the lake. We estimated that the Tilapia farm was responsible for ~ 86% of nitrogen (N) and ~95% of the phosphorus (P) annual contributions to Lake Yojoa. This disproportionate nutrient loading of both N and P suggests that this single aquaculture operation, more so than changes in nutrients inputs from the watershed, was responsible for the previously documented deterioration of the Lake Yojoa ecosystem. This study shows the potential for net-pen aquaculture to have disproportionately negative impacts on freshwater ecosystems, even when operations meet all standards for the current state-of-the-art sustainability certifications. We suggest shifts in metrics that could improve the impact of the certification process so that best practices can reduce the impact of net-pen aquaculture on freshwater ecosystems and arrive at the intended goal of long-term environmental sustainability.

## Introduction

The majority of nutrients in inland waters come directly from terrestrial surfaces such as agricultural run-off and untreated municipal effluent which can alter in-stream nutrient concentrations (Tromboni and Dodds 2017, Gücker et al. 2006), stimulate autotrophic and heterotrophic microbial metabolism (Sterner and Elser 2003, Dodds 2006), and increase the transport of reactive nutrients downstream, often altering trophic state within the downstream ecosystems. While terrestrial inputs dominate most aquatic nutrient budgets, net-pen aquaculture, a well-established production technique used in the rapidly growing finfish rearing industry (Naylor et al. 2021), can also contribute a substantial amount of nutrients to the systems they inhabit. Though net-pen production occurs across nearly all latitudes, many inland net-pen operations are in tropical regions (FAO 2022). Understanding the sources and impacts of nutrient loading to aquatic ecosystems requires *in situ* monitoring and access to administrative and fiscal support. Unfortunately, many countries in tropical regions lack the resources to study and monitor inland waters at the same scale as countries in the ‘Global North’, leading to an unequal distribution of ecological data across latitudes (Finkler et al. 2018, Riveros-Iregui et al. 2018). In addition to limitations of extant data, the surrounding terrain in tropical regions can be harder to physically access (private land ownerships) and/or may have rugged or undeveloped access (e.g., irregularly graded roads that require 4×4 capability).

Quantitative estimates of watershed scale loading of nutrients and quantified tributary contributions to lake ecosystems are rare in temperate zones and largely restricted to larger lakes (Robertson and Saad 2011, North et al. 2013, Mooney et al. 2020) and are even rarer in the tropics and subtropics (Coveney et al. 2005, Li et al. 2011, Luo et al. 2022). Whereas many North American streams and rivers have been instrumented by the United States Geological Survey (USGS), no comparable effort to monitor discharge has yet to be broadly implemented for lower latitudes (Riveros-Iregui et al. 2018). This deficit of discharge data and the absence of direct measurements or sufficient data to employ modeling techniques, such as SPAtially Referenced Regressions on Watershed attributes (SPARROW) models (Preston et al. 2011), makes assessing the relative impact of various point and non-point sources of nutrients difficult if not impossible. Many of these challenges can be addressed through local collaborations, highlighting the importance of relationships with regional non-profit organizations and extant, although often limited, regional research programs. Such organizations provide critical local knowledge, such as seasonality of discharge or other ecosystem characteristics, that aids in experimental design, a better understanding of catchment boundaries, and how to access specific points within catchment areas for optimal monitoring.

The primary objective of this study was to identify the dominant sources of nutrients to a large tropical lake which has seen rapid eutrophication in recent decades (Fadum and Hall 2022). We compared the nutrient contributions of the six main tributaries that feed Lake Yojoa (Honduras) to total annual nutrient inputs from the large Tilapia operation that is located in the north central region of the lake (Figure 1). To do this we 1) characterized the seasonality of discharge and geochemistry in Lake Yojoa’s main tributaries, 2) quantified annual watershed nutrient contributions to Lake Yojoa, and 3) estimated the contributions of nutrients supplied to Lake Yojoa by a large aquaculture operation.

**Figure 1.**
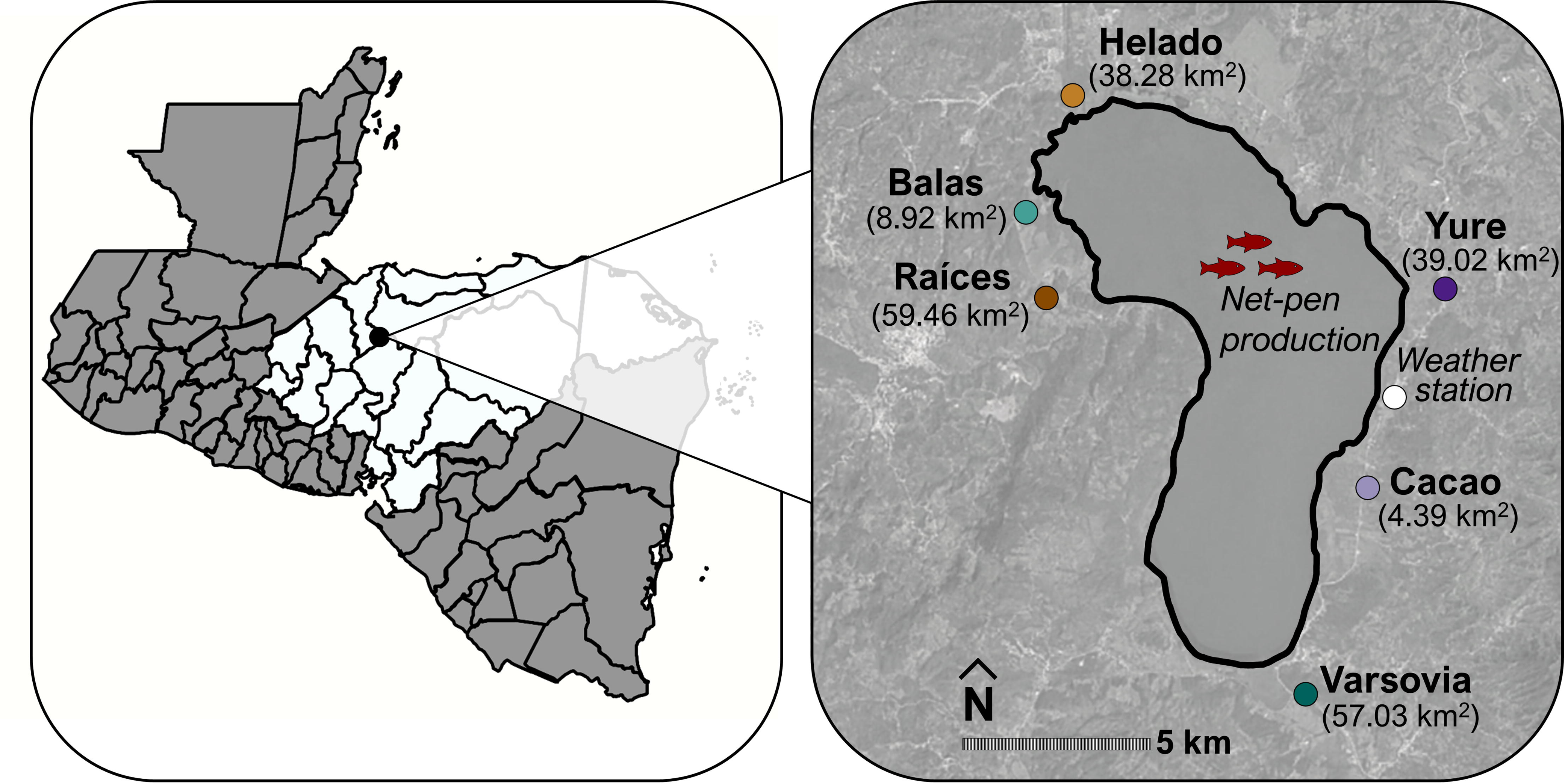
Map of the Yojoa watershed, identifying the six tributary sampling points (with drainage area in parentheses) as well as the weather station and aquaculture operation locations. Country outlines were generated using MapChart (www.mapchart.net, 2023; Creative Commons Attribution-ShareAlike 4.0 International License). Aerial imagery from Copernicus, accessed through Google Earth Pro.

## Methods

### Study location

Lake Yojoa (83 km^2^) is located in the neotropics of Honduras and is the country’s largest freshwater lake. The lake and its surrounding watershed (440 km^2^, approximately ~200 km^2^ of which directly contributes to Lake Yojoa) supports nine municipalities and a variety of land use/land covers (LULC). The region is marked by seasonal monsoonal precipitation (usually from late June – October) resulting in high volume and “flashy” riverine inputs. Lake Yojoa is located at the center of the watershed with six principal contributing tributaries (Balas, Cacao, Raíces, Helado, Varsovia, and Yure) (Figure 1).

Sampling sites were selected based on stream size and potential relative influence on overall water quality to Lake Yojoa (Figure 1). All stream sampling sites were chosen to be as close to the lake inflow location as possible while also selecting a reach where stream geomorphology was constrained and unlikely to change during the course of the project. This resulted in sampling locations with consistent channel morphology, stable banks, and bottom structure (overpasses with cement structure were used as often as possible), and minimal morphology change for the best estimation of discharge measurements.

### Discharge measurements

Rating curves were constructed following USGS methodology for cross-sectional depth and velocity measurements at various flows (Dingman 2002, Supplemental Text 1, Figure S1). In four principal tributaries of Lake Yojoa (Balas, Helado, Raíces, and Varsovia), we deployed HOBO U20L-01 pressure transducers set at 15-minute intervals. We installed pressure transducers in compliance with EPA Protocols for Measuring Water Level and Streamflow (Meals and Dressing 2008). Pressure transducers tracked water levels which, when coupled with ratings curves, allowed for estimation of discharge at 15-minute intervals over the duration of our monitoring efforts (April 2019-April 2020). Staff gauges were also installed at each sampling location for visual confirmation of stage height and pressure transducer derived discharge measurements were corrected using weekly staff measurements (Supplemental Text 1, Figure S2). See data availability statement for code used to calculate discharge in Balas, Raíces, Helado, and Varsovia.

Where pressure transducer installation was not feasible (Cacao and Yure), we used Manning’s equation (Equation 1) to measure discharge at the channelized locations. The channel slopes for Cacao and Yure were 0.0002 and 0.002, respectively. We used Manning’s roughness coefficient of 0.013 (Chow 1959).

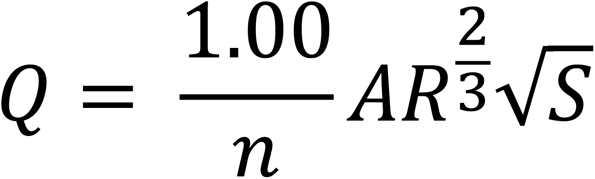

**Equation 1**. Manning’s Equation to estimate discharge in tributaries Yure and Cacao. n= Manning’s roughness coefficient, A= Flow area (ft^2^), R= Hydraulic radius (ft), S= Channel slope

### Assessing the impact of sampling frequency

To address the impact of sampling frequency on estimates of daily mean discharge, we tested how infrequently discharge could be measured to identify a mean discharge value that was within 10 percent of the most accurate estimate of annual discharge (i.e., modeled value based on the measurement of stage at 15-minute intervals) in our four ‘high-frequency’ tributaries (Raíces, Balas, Helado, and Varsovia). We tested the following scenarios: 1) apply the observation taken on the first day of each month to the whole month (i.e. sampling frequency = 12), 2) use observations collected twice a month (i.e. sampling frequency = 24), 3) weekly observations (i.e. sampling frequency = 52), 4) daily observations (i.e. sampling frequency = 365), 5) measure discharge twice daily at 8:00 and 16:00 (i.e. sampling frequency = 730), 6) measure discharge three times daily at 6:00, 12:00, and 18:00 (i.e. sampling frequency = 1095), 7) measure discharge four times daily at 5:00, 10:00, 15:00 and 20:00 (i.e. sampling frequency = 1460), and 8) measure discharge hourly (i.e. sampling frequency = 8760). While the minimum sample frequency varied between tributaries and across months (Supplemental Text 2, Figure S3 and S4), it demonstrated that high temporal resolution (i.e. 15-minute intervals) was not necessary for accurately assessing discharge. Particularly in the dry season, weekly measurements were typically sufficient to accurately (i.e., estimating discharge within +/- 10% of discharge estimated hourly) assess discharge. Therefore, we believe that the lower sampling frequency in the tributaries we were unable to instrument did not decrease the accuracy of the estimates of discharge or our estimates of load. We also further corrected for any potential methodological differences in how sampling frequency impacted our calculations of nutrient loading, as described below.

### Sample collection and analyses

In collaboration with Asociación de Municipios del Lago de Yojoa y su Área de Influencia, (AMUPROLAGO) we collected weekly water samples in duplicate at all six tributary locations. Water samples were collected in acid washed 50 ml conical tubes (Falcon 352070) and immediately placed in a cooler on ice for transport back to AMUPROLAGO. Water samples were then frozen at 4 °C until transported back to Fort Collins, CO, USA on ice. All analyses were performed at the EcoCore Laboratory at Colorado State University (CSU, Fort Collins, CO, USA). All water samples were analyzed for total phosphorous (TP), dissolved organic carbon (DOC), total dissolved nitrogen (TDN), ammonium (NH_4_^+^-N, hereafter referred to as NH_4_^+^), and nitrate (NO_3_^−^-N, hereafter referred to as NO_3_^−^). A complete description of methods of chemical analyses can be found in previous publications (Fadum and Hall 2022, Fadum et al. 2023, Fadum et al. 2024).

### Nutrient loading estimates

The six tributaries in this study exhibited largely chemostatic behavior. Chemostatic systems are defined by the absence of a consistent relationship between discharge and the concentration of the analyte of interest (i.e., a slope of zero in log-log space, Godsey et al. 2009, Wymore et al. 2017). Chemostasis means it was not possible to estimate nutrient concentration from discharge measurements because of the lack of consistent statistically significant relationships between nutrient concentration and discharge across all months in the six tributaries (Discussed in Results below, Table 1, Figure 4). Therefore, in tributaries with daily discharge measurements (Balas, Helado, Raíces, and Varsovia), nutrient concentrations were interpolated between weekly nutrient measurements (assuming linear drift) to approximate daily nutrient concentrations and then multiplied by daily mean discharge to calculate daily nutrient loading. This method is consistent with well-established field methods for estimating nutrient flux (Johnson et al. 1969).

**Table 1.**
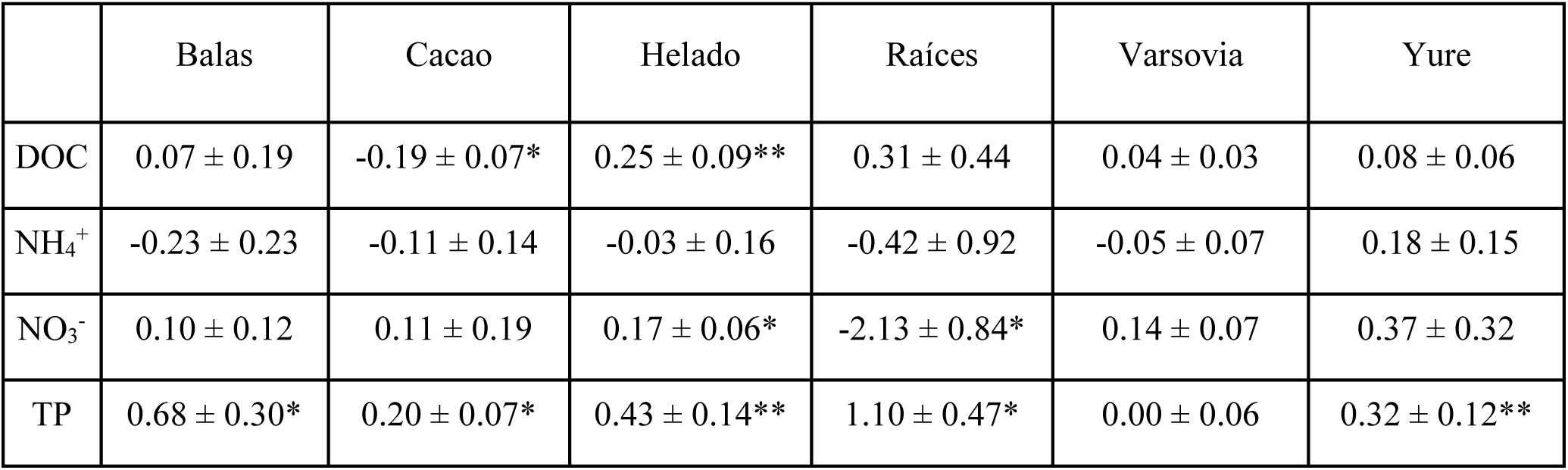
Slope (±SE) of the relationship between in-tributary nutrient concentrations and discharge in log-log space and significance. A chemostatic relationship is defined at a slope of zero. Significance codes: ‘***’ 0.001 ‘**’ 0.01 ‘*’ 0.05

In the two tributaries where Manning’s equation was used (Cacao and Yure), interpolation to arrive at daily estimates of load was not possible because we had only weekly measurements of discharge which matched our weekly measurements of nutrient concentrations. To account for the temporal differences in loading estimates between Cacao and Yure and the other four principal tributaries, annual cumulative loading estimates for each tributary were normalized by the number of available daily loading values. The sum of these additional inputs, which are not reflected in the comparison of monthly tributary inputs, is identified as ‘tributary underestimation’ in the comparison of tributary and aquaculture inputs. Therefore, while we may have underestimated the contributions of Cacao and Yure individually, we have likely accurately accounted for the overall nutrient contributions from the six principal tributaries to the lake.

Estimates of aquaculture inputs for the year 2013 came from Regal Springs Aquaculture LLC as part of a collaborative data sharing agreement between AMUPROLAGO and CSU. Total N and P loading was determined by N and P content of feed provided less N and P recovered in harvested biomass. Because the number of net pens in Lake Yojoa have increased from ~78 in 2013 to ~163 in 2020, we consider these values as conservative estimates of aquaculture nutrient loading for the 2019-2020 comparison. The probable corresponding increases in contemporary nutrient contributions are estimated and further described in the discussion section (Supplemental Text 3, Figure S5-S7).

### Analysis and calculations

The calculations described above were conducted in R (version 4.2.2). Figures were similarly constructed in R, using *ggplot2* (Wickham 2009). Discharge correlation coefficients were calculated using *psych* (Revelle 2024). For a full list of packages used, as well as details of data analysis, see provided code repository in the Data Availability Statement. Drainage areas were calculated using ArcGIS Pro’s watershed tool at 90-meter resolution.

## Results

Between April of 2019 and April of 2020, The Lake Yojoa watershed experienced ~5-6 months with limited precipitation (November-April) and ~6-7 months with pronounced precipitation (April-October) with most of the annual precipitation falling between July and October (Figure 2). This is consistent with previously described patterns of seasonal precipitation for the Yojoa watershed (Fadum and Hall 2022). These seasonal differences in precipitation result in pronounced differences in delivery of reactive nutrients from the watershed to Lake Yojoa among seasons. Here we first show the differences in discharge, nutrient concentration, and loading for the six principal tributaries that transport nutrients from the Lake Yojoa Watershed to Lake Yojoa and then we compare those inputs to our best estimates for inputs from the industrial aquaculture operation in Lake Yojoa.

**Figure 2.**
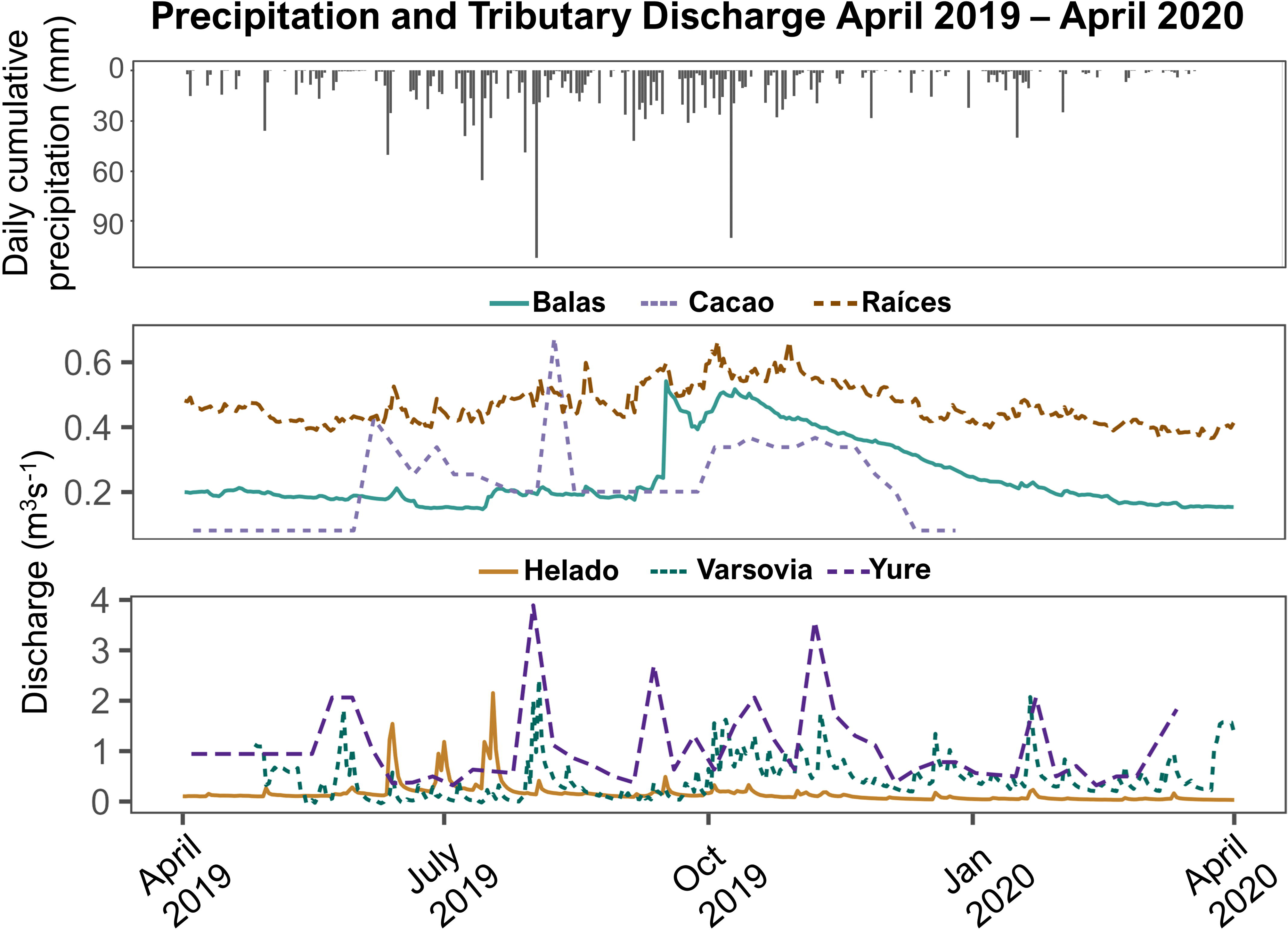
Hyetograph (top panel) with discharge for duration of study (April 2019-April 2020) for low discharge tributaries Balas, Raíces and Cacao (middle panel) and high discharge tributaries, Yure, Varsovia and Helado (bottom panel).

### Discharge

We first evaluated the discharge from each of the six principal tributaries to Lake Yojoa. Three (Balas, Cacao and Raíces) of the six tributaries experienced persistently low (< 1 m^3^s^-1^) discharge throughout the monitoring period (April 2019-April 2020). Of the three low-flow tributaries, Raíces had the least amount of seasonal variation, and exhibited low and relatively consistent discharge (min-max, mean ± SE; 0.31-0.68, 0.45 ± 0.00 m^3^s^-1^, Figure 2). Conversely, Cacao’s discharge was more closely coupled to rainfall events and therefore “flashier”. Cacao also supported minimal baseflow during the dry months and peak discharge was concomitant with the rainy season (0.08-0.67, 0.18 ± 0.01 m^3^s^-1^, Figure 2). Discharge in Balas also peaked during the rainy season and decreased during the dry months (0.12-0.54, 0.23 ± 0.00 m^3^s^-1^, Figure 2). However, unlike Cacao, Balas’s hydrograph suggested that its discharge was connected to groundwater inputs, due to the threshold like peak response and gradual decline of that discharge throughout the dry season (Figure 2). This is consistent with the origin of Balas (~200 m upstream from where we measured discharge) being from a mountainside.

The remaining three tributaries (Helado, Varsovia, and Yure) experienced episodic flows which exceeded the maximums of the previously discussed tributaries. Helado experienced peaks during the rainy season concomitant with sustained large rain events (0.02-2.15, 0.13 ± 0.00 m^3^s^-1^, Figure 2). The hydrology of Yure (0.32-3.89, 1.07 ± 0.11 m^3^s^-1^, Figure 2) and Varsovia (0.00-2.56, 0.51 ± 0.02 m^3^s^-1^, Figure 2) were decoupled from seasonality of precipitation as they were sustained by reservoir release and thus did not respond as directly to individual precipitation events. In addition, Yure and Varsovia are cement channelized tributaries, thus preventing groundwater seeps.

Despite differences in the hydrographs of the six principal tributaries, there were moderate correlations between the discharge measurements in the four tributaries where we were able to measure daily mean discharge (Balas, Raíces, Helado, and Varsovia). Balas and Raíces, which are co-located in the northwest region of the watershed, had the most highly correlated discharge, with a positive correlation (r=0.82) across all seasons. Discharge in Raíces also showed a weak but positive correlation (r=0.53) with discharge from Cacao as well as Varsovia (r=0.42). Similarly, Varsovia and Balas had a weak correlation between their discharge (r=0.46).

### Tributary nutrient concentration

Water-column, nutrient concentrations differed among the six principal tributaries and among the three principal elements we tracked, C, N and P.

#### Phosphorus

In-tributary TP concentrations in all six tributaries were generally below 3 µM with the exception of some intermittent peaks in Raíces, Varsovia and Yure. Balas had the lowest in-tributary TP concentrations (min-max, mean ± SE; 0.09-1.61, 0.48 ± 0.04m^3^ µM, Figure 3A). Despite occasional peaks, Yure (0.16-2.71, 0.52 ± 0.04 µM, Figure 3A), Varsovia (0.12-3.13, 0.66 ± 0.05 µM, Figure 3A) and Helado (0.09-2.22, 0.61 ± 0.04 µM, Figure 3A) had similarly low mean TP throughout the monitoring period (April 2019-April 2020). Cacao had a moderately higher mean annual TP concentration (0.48-2.06, 0.83 ± 0.03 µM, Figure 3A). Raíces (0.25-4.93, 1.30 ± 0.07 µM, Figure 3A) had the highest concentrations of TP. Despite the predominantly chemostatic behavior across the six tributaries (Figure 4), we did observe weak but significant and positive relationships between TP concentrations and discharge measurements in all tributaries with the exception of Varsovia (Table 1).

**Figure 3.**
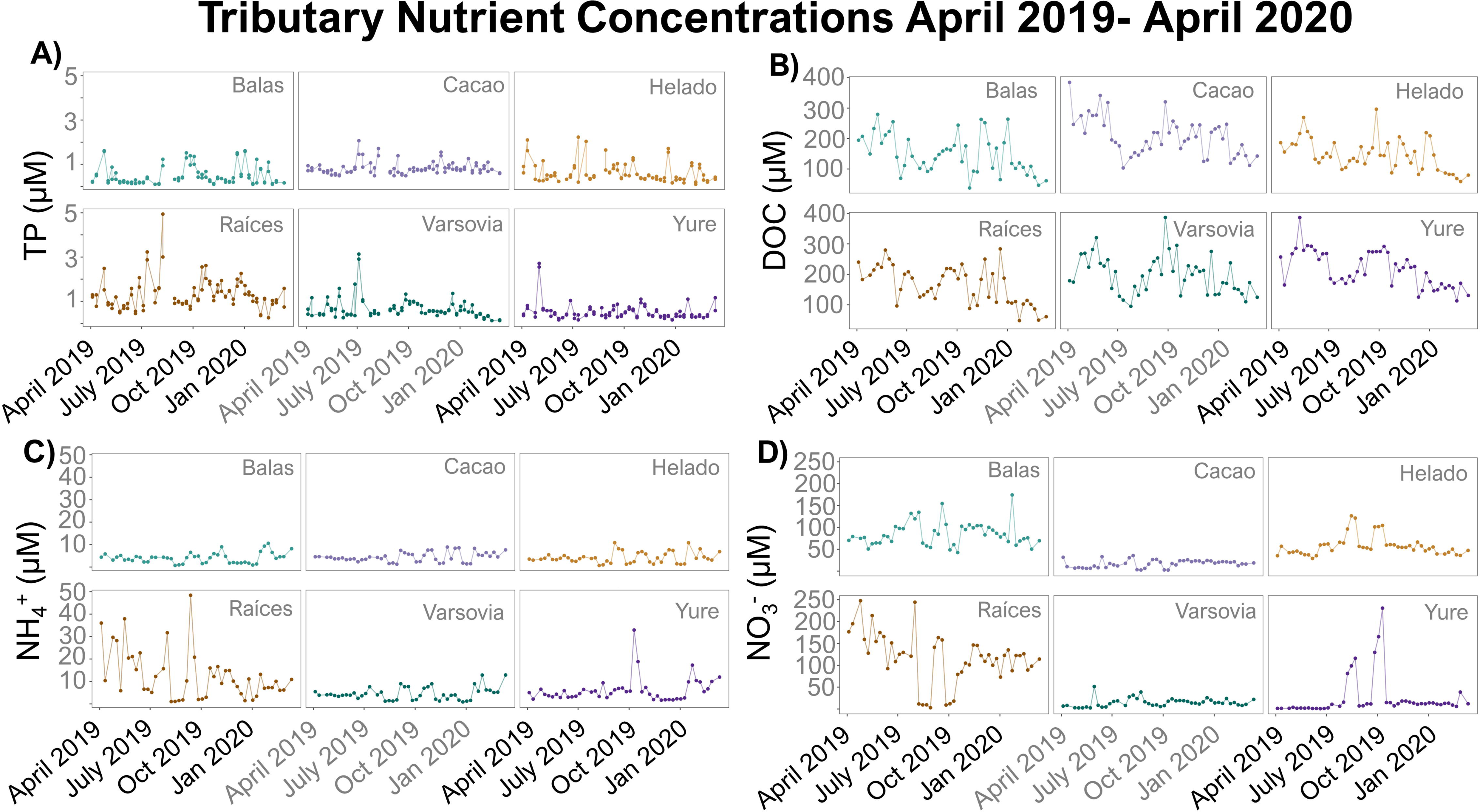
In-tributary nutrient concentrations of **A)** TP, **B)** DOC, **C)** NH_4_^+^, and **d)** NO_3_^−^ in all six tributaries.

**Figure 4.**
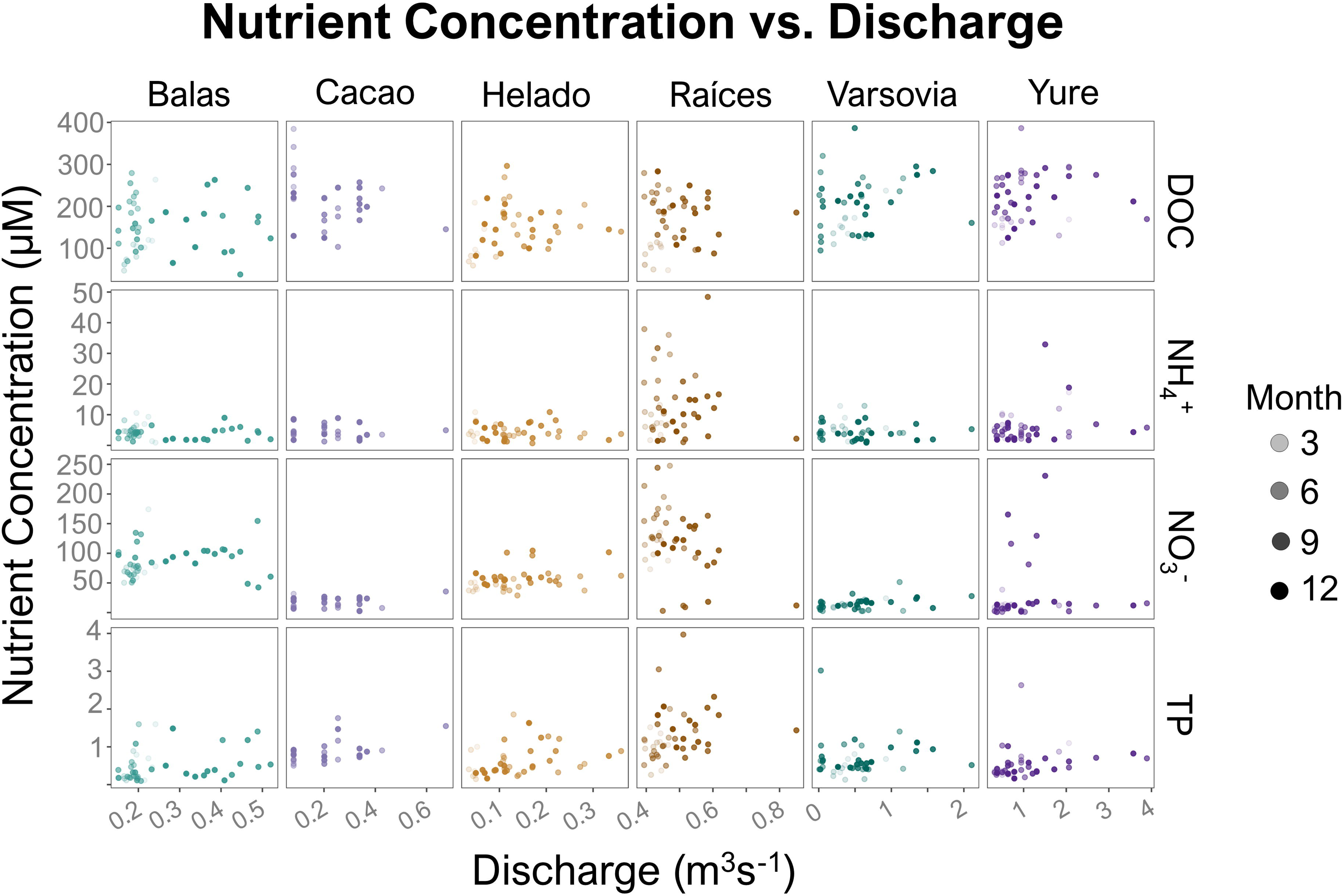
In-tributary nutrient concentrations versus discharge in all six tributaries. Slope in log-log space provided in Table 1.

#### Carbon

All six tributaries exhibited similar patterns in DOC. In Balas (min-max, mean ± SE; 19.56-325.03, 150.86 ± 7.86 µM, Figure 3B), Helado (43.46-371.82, 148.86 ± 6.72 µM, Figure 3B), Raíces (43.62-314.12, 165.93 ± 7.11 µM, Figure 3B), Cacao (98.24-443.59, 209.19 ± 8.05 µM, Figure 3B), Varsovia (86.58-405.46, 199.66 ± 7.46 µM, Figure 3B), and Yure (101.90-481.34, 218.82 ± 6.79 µM, Figure 3B) we observed a decrease in DOC concentrations during the rainy season followed by an increase in DOC entering the dry season. Only Cacao and Helado (the two tributaries that responded most to episodic rainfall, Figure 2) had significant relationships between DOC concentrations and discharge (Table 1).

#### Nitrogen

In-tributary concentrations of NH_4_ ^+^ were generally low with the exception of Raíces (min-max, mean ± SE; 0.92-49.19, 13.03 ± 1.15 µM, Figure 3C) which had greater seasonal variation in observed NH_4_ ^+^ concentrations than the other five tributaries. The remaining tributaries had consistently low NH_4_ ^+^ concentrations. Balas (0.85-13.06, 4.20 ± 0.26 µM, Figure 3C), Cacao (1.07-12.27, 4.60 ± 0.25 µM, Figure 3C), Helado (0.64 - 14.06, 4.31 ± 0.30 µM, Figure 3C) and Varsovia (0.57 - 16.84, 4.73 ± 0.32 µM, Figure 3C) were similar across seasons. Yure’s otherwise low NH_4_^+^ concentrations were punctuated by occasional peaks (0.71 - 36.05, 6.00 ± 0.60 µM, Figure 3C). Despite the episodic peaks in NH_4_^+^, we observed no significant relationships in any of the tributaries between NH_4_^+^ concentrations and discharge (Figure 4, Table 1).

The behavior of NO_3_^−^ was different among each of the tributaries. As with NH_4_^+^, Raíces had the greatest concentrations of NO_3_^−^ and highest variance in concentration throughout our study (min-max, mean ± SE; 2.85-254.66, 120.54 ± 5.87 µM, Figure 3D). Balas had lower mean NO_3_^−^ and a less pronounced variation than Raíces, (10.06-174.20, 86.07 ± 3.06 µM, Figure 3D). Helado (21.27-105.66, 54.96 ± 1.92 µM, Figure 3D) and Yure (0.07-248.30, 24.21 ± 4.88 µM, Figure 3D) had fewer peak events compared to Balas and Raíces, resulting in a lower annual means. Varsovia (1.49-52.04, 14.10 ± 0.98 µM, Figure 3D) and Cacao (2.21-43.12, 17.61 ± 0.95 µM, Figure 3D) had least variation in NO_3_^−^ concentrations and the lowest mean NO_3_^−^. However, both Helado and Raíces did have significant correlations between discharge and NO_3_^−^ concentrations (Table 1).

### Tributary vs. Aquaculture loading

These differences and dynamics in stream nutrient transport led to variation among months and among tributaries in the amount of reactive nutrients contributed to Lake Yojoa by the six main tributaries (Figure 5A-B). When the tributary contributions were combined (i.e. summed for each month), October had the most pronounced contribution for both TP and TDN (Figure 5C-D). Raíces was the top contributor of both TDN (28,432 kg) and TP (643 kg) (Table 2). Balas was the second greatest contributor of TDN (10,088 kg). Varsovia was the second greatest contributor of TP (275.50 kg) (Table 2).

**Figure 5.**
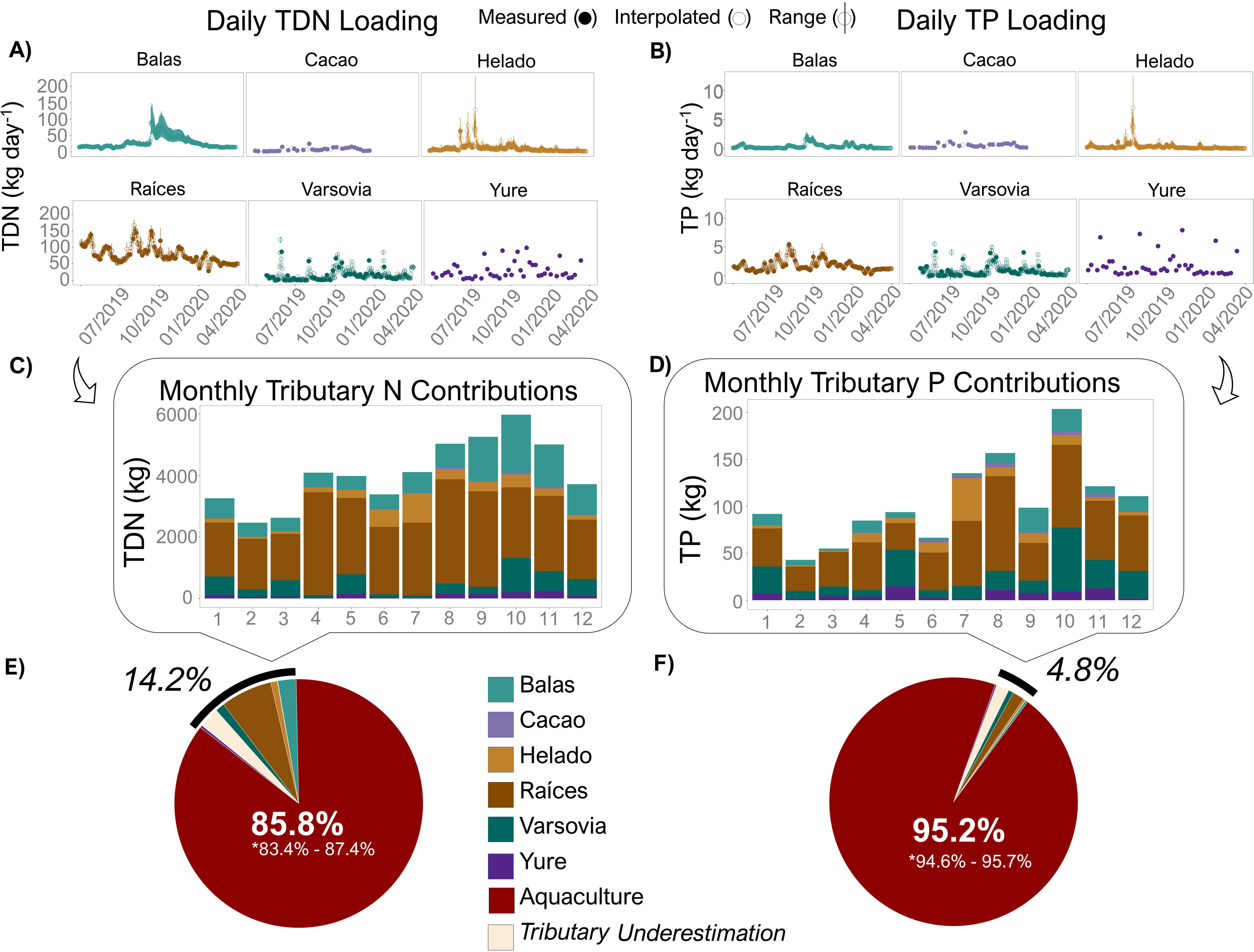
Loading of **A)** TDN and **B)** TP, open circles are days where estimated loading has been interpolated. The cumulative normalized difference is provided in an 8^th^ category in panels E and F. Monthly tributary loading for **C)** TDN, and **D)** TP in the six tributaries. **E)** Annual total tributary N contributions compared to aquaculture estimated loading *with range reflecting minimum and maximum modeled discharge. **F)** Annual total tributary P contributions compared to aquaculture estimated loading *with range reflecting minimum and maximum modeled discharge.

**Table 2.** Annual nutrient loading for Lake Yojoa’s major tributaries and industrial net-pen aquaculture.

| Source | TDN loading (kg) | TDN per area (kg km<sup>-2</sup> or kg per net-pen) | TP loading (kg) | TP per area (kg km<sup>-2</sup> or kg per net-pen) |
| --- | --- | --- | --- | --- |
| Cacao | 276.4 | 62.8 | 20.6 | 4.6 |
| Yure | 1,320.2 | 33.8 | 79.1 | 2.0 |
| Helado | 3,638.9 | 95.0 | 114.6 | 2.9 |
| Balas | 10,088.3 | 1,129.9 | 129.2 | 14.4 |
| Varsovia | 5,138.0 | 90.0 | 275.5 | 4.8 |
| Raíces | 28,432.8 | 478.1 | 643.2 | 10.8 |
| Aquaculture | 364,052.3 | 4,667.3 | 39,768.6 | 509.8 |

However, the tributaries were responsible for only a small fraction of the estimated nutrient load to Lake Yojoa. Using the calculated nutrient loading provided by the aquaculture operators, we estimated that their aquaculture operation contributed 364,052 kg N and 39,768 kg P during a single calendar year (Supplemental Text 4, Table S4). When we compared the annual sum of watershed loading as described above to the best estimates for nutrients derived from the aquaculture operation, we calculated that aquaculture was responsible for 95.2% of annual TP loading and 85.8 % of the annual nitrogen loading to Lake Yojoa (Figure 5E-F).

## Discussion

This study represents the first known comparison of the relative impacts of nutrient loading between a large (440 km^2^) multi-use watershed and a sustainably certified aquaculture operation to a large, tropical lake (83 km^2^). This comparison illustrates the disproportionate impact that large aquaculture operations, even those using best practices and meeting the highest industry standards for minimizing ecosystem impact, can have on freshwater ecosystems. Here we discuss the findings of our analyses, the challenges in conducting research on the impacts of aquaculture, and suggest potential paths forward to create better practices in sustainable net-pen aquaculture in order to maintain the integrity of freshwater ecosystems for the foreseeable future.

### Nutrient Loading from a Large Tropical Watershed

Our estimates likely captured the majority of dissolved nutrient inputs to Lake Yojoa from the surrounding watershed. Watershed monitoring in Latin America (and tropical latitudes in general) pales in comparison to watershed monitoring in temperate latitudes across the globe (Riveros-Iregui et al. 2018). Within this watershed analysis, we found that Raíces was the largest transporter of reactive nutrients to Lake Yojoa. Raíces is a relatively wide river capable of massive, if irregular, flows, but with consistent, relatively low base flow throughout the year. The Raíces sub-watershed contains the largest municipality in the Lake Yojoa basin, Las Vegas (est. pop. ~28,000, Instituto Nacional de Estadística Honduras), as well as significant agricultural and mining activities resulting in a water column that is enriched in reactive nutrients compared to the water columns of the other principal tributaries (Figure 3).

In two of the tributaries (Yure, cement channel draining a small reservoir, and Cacao, a small tributary draining a relatively protected sub-watershed that originates within a national park, Parque Nacional Cero Azul Meámbar) we were unable to install pressure transducers and thus unable to estimate daily flows. However, because the entirety of Yure was cement channelized and the portion of Cacao where we measured discharge and collected nutrient samples was also cement channelized, we were able to use weekly measurements of stage and Manning’s equation to estimate discharge. This created the added challenge of having four tributaries with high temporal resolution estimates of stage (15-minute estimates of stage, averaged to daily discharge) and two tributaries with lower temporal resolution (weekly measurement of stage, interpolated to daily discharge) from which to calculate discharge and thus total annual loading. We addressed these differences in two ways; first by assessing minimum sampling frequency required to approach our best estimates of annual load from the Yojoa watershed (Supplemental Text 2), and second by estimating the cumulative loading from days which we were unable to interpolate (‘Tributary underestimation” in Figure 5E and F). These calculations provide us with further confidence that we have accurately estimated the contributions of Yure and Cacao and we have determined an accurate if conservative estimate of nutrient contributions from each tributary to Lake Yojoa.

To further assess our estimates of annual watershed load relative to the aquaculture contributions, we also recalculated watershed contributions using the maximum modeled discharge (instead of median modeled discharged). When we did this (following all the same methods detailed above) the increase in watershed nutrient contributions remained inconsequential compared to the nutrient contributions of aquaculture, reducing estimated contributions of P from 95.2% to 94.6% and contributions of N from 85.8% to 84.4%.

In addition to sampling frequency, we assessed additional potential sources of watershed nutrient loading underestimation. First, we evaluated the potential impact of comparing total N (aquaculture additions) to dissolved N (tributaries, as total N measurements were unavailable). When we compared particulate N (size fraction ≥ 0.7 μm) to total dissolved N (size fraction ≤ 0.7 μm) in tributaries in June 2019, we found the majority of tributary N to be in the dissolved fraction (Supplemental Text 5, Table S5). Global estimates of in-river NO_3_^−^ to total N suggest a 1:1 ratio of dissolved to total N (Turner at al. 2003). Therefore, we can reasonably expect Lake Yojoa Watershed N contributions to exceed loading determined by the dissolved fraction alone. However, even if we were to double our estimations of watershed N contribution to account for this potential under-sampling, aquaculture would still account for 75% (instead of 85.8%) of N contributions to Lake Yojoa.

We also assessed potential contributions of atmospheric deposition. Regional estimates of N deposition for the area of Lake Yojoa are approximately equivalent to the annual contributions of Varsovia (5922 to 7644 kg yr^-1^, Dentener 2006, Fadum and Hall 2022). Therefore, whereas atmospheric deposition likely contributed to Lake Yojoa’s annual nutrient load, best published estimates for this region suggest it would not be a sizeable enough fraction to alter the relative impact of aquaculture nutrient additions. Finally, all six tributaries pass through retention ponds or wetlands before entering Lake Yojoa and thus our estimates of nutrient loading (taken upstream of the integration with the lake) likely overestimate delivery of reactive nutrients from each sub-watershed. Taken together, these points suggest our estimates of reactive N and P loading from the Lake Yojoa watershed to Lake Yojoa to be accurate if perhaps a bit conservative. This is especially true if we are only comparing the relative contribution of watershed nutrients to aquaculture derived nutrients.

### Nutrient Loading from An Industrial Net-Pen Tilapia Operation

In general, nutrient loading data from aquaculture companies are not publicly available and, even though required for various certifications, rarely published. However, due to a data sharing agreement in 2013, the operators of the industrial aquaculture operation shared detailed monthly estimates of mass of monthly feed, percent N and P of feed, wet biomass of Tilapia harvest, and percent N and P of harvested Tilapia (Supplemental Text 4). Using these detailed data, we estimated the N and P loading from aquaculture to Lake Yojoa for 2013, the most complete annual data provided. Analyses of images from Google Earth clearly show that in 2013 the industrial aquaculture operation maintained 78 net pens on Lake Yojoa. In January of 2019 the same company maintained ~148 net pens on the lake and in May of 2020 that number had increased to 163 net pens on Lake Yojoa (Supplemental Text 3, Figure S5). If we assume that nutrient loading scales linearly with the number of net-pens, it is likely that contemporary inputs (those from 2019 and 2020) from aquaculture was ~ 48% greater than the loading values presented here (Figures S4 and S5). The 2013 loading from the aquaculture operation was already ~ 86% of nitrogen (N) and ~95% of the phosphorus (P) contributions to Lake Yojoa annually and has likely dramatically increased since then. The overwhelming contribution of the aquaculture activities to the annual nutrient inputs to Lake Yojoa suggests that the documented eutrophication of Lake Yojoa over the past 40 years (Fadum and Hall 2022) is almost entirely due to nutrient inputs from the aquaculture operation within Lake Yojoa.

### Challenges with Current Sustainability Certification Practices and Suggestions for Improving Current Standards

Despite the industrial aquaculture operator’s obtaining multiple top sustainability standards (as of 2023), their nutrient contributions have unequivocally contributed to the rapid ecosystem deterioration observed in Lake Yojoa in recent decades (Fadum and Hall 2022). This highlights the ways in which the monitoring required for sustainability certification for net-pen aquaculture is maladapted to the ecosystems hosting the aquaculture operations or else fall short of sufficiently preventing environmental degradation (Boyd et al. 2005, Rector et al. 2023). As such, certifying agencies are not only failing to deliver on their promise of environmental stewardship and failing to protect the interests of local communities and the investments made by aquaculture farm operators, but they are also creating a production pipeline that misrepresents that product to the consumer. Below we identify ways in which current sustainability standards specific to net-pen aquaculture could be improved towards the goal of increased ecological relevance.

Lake Yojoa’s large aquaculture operation is accredited by multiple independent third-party certifiers including the Aquaculture Stewardship Council (ASC). Whereas the entire certification process is too detailed for description in this manuscript, briefly, the ASC standards for net-pen reared Tilapia require the operator to collect monthly data Secchi depth and dissolved oxygen, turbidity, specific conductance, algal density (measured as Chlorophyll *a* μg L^-1^), phosphate and ammonia from surface waters (1m). Samples are collected at three locations: 1) a ‘Receiving Water-Reference Point’ (RWRP), defined as the point within a lake at the “maximum distance from the influence from the farming activities”, 2) a ‘Receiving Water-Farm outfall or mixing zone’ (RWFO), a point within the cages, and 3) ‘Receiving Water-Farm afar’ (RWFA), defined as a point “down the prevailing current pattern in a lake” (ASC Tilapia standards v1.2). Measurements between each location are compared to assess long term impacts. However, current ASC standards only require that operators maintain ≤ 65% change in diurnal dissolved oxygen, ≤ 20 μg L^-1^ P in and ≤ 4 μg L^-1^ Chlorophyll *a* in surface waters, and limit annual P loading to 20 kg per metric ton of Tilapia produced. An express limit on P only without consideration of N may be particularly problematic given the varied and heterogenous nutrient limitation landscape of tropical lakes (Fadum and Hall 2023).

From an ecosystem perspective, these criteria to measure ecosystem impact create several challenges. Principal among these is that water columns of deep (>12m) lakes are often seasonally stratified and surface water chemistry is likely to be very different from deep water chemistry. This is further complicated by algal production and fish waste not being neutrally buoyant and therefore passively exported from the surface waters to the deep waters. Thus, monitoring of surface waters is likely to miss the principal impacts of aquaculture on lake ecosystems by failing to capture nutrient export via algal biomass and fish waste from the surface to the deep water. Accumulation of excessive nutrients in the deep water can result in high surface nutrient content, as previously observed in Lake Yojoa (Fadum and Hall 2022). This deep water nutrient accumulation can temporally decouple the impact of aquaculture nutrient input and ecosystem response because the nutrient inputs can accumulate outside the photic zone for a large part of the year. During annual turnover (when bottom waters mix with surface waters) the abundance of accumulated nutrients in the aphotic deep water will stimulate phototrophy and primary production in the surface waters. Thus, algal blooms occur episodically with annual (and other smaller intra-annual) water column inversions separating the daily inputs of aquaculture derived nutrients from the episodic algal productivity events that they fuel. The increase in frequency and intensity of algal blooms, as observed in Lake Yojoa, leads to gradual alteration of ecosystem state (Fadum and Hall 2022). To improve current aquaculture certification standards, nutrient concentrations of the surface and bottom layer must both be considered in order to make sure there is not cumulative change in either with the potential to alter ecosystem state during turnover events.

A second challenge with the current metrics for certification is the comparison between reference sites (e.g., RWRP) and impacted (RWFO and RWFA) sites. Physical dynamics of lake ecosystems are complicated (Imberger 1998, Bruce et al. 2018, Amorim et al. 2021) and it is rare to have a fully described three or even two-dimensional understanding of water column dynamics. Thus, having a single or a second reference point is likely insufficient to characterize how nutrient inputs are impacting a whole lake ecosystem. To better address impacts of aquaculture on whole lake ecosystems, aquaculture certifications should require a more spatially sufficient coverage to better characterize the entire ecosystem. This approach would increase ecosystem coverage of nutrient concentrations, algal abundance, and turbidity, and if designed appropriately could be coupled with satellite overpasses to create a model with the potential for passive monitoring of ecosystem state through remote sensing (Dörnhöfer and Oppelt 2016). This would improve both spatial coverage and temporal resolution of ecosystem monitoring over time and help monitor the trophic state of the entire ecosystem.

A third challenge to understanding the impact of aquaculture on lake ecosystems is understanding the nutrient buffering capacity of lake sediments. Reactive nutrients exported from surface water as particles often settle into reactive anoxic sediments. When the water column is mixed, there is a sharp oxic/anoxic gradient at the sediment water interface creating the potential for a biogeochemical “hotspot” (McClain et al. 2003). For example, when the water column is mixed, degradation of organic matter within the sediments can lead to a steady diffusion of NH_4_^+^ to the water column. This sediment reservoir may continue to supply a lake with NH_4_^+^ even when monitored water column nutrient concentrations have decreased, resulting in a delay between decreased nutrient loading and ecosystem recovery, as has been observed in Lake Yojoa (Fadum et al. 2023). Thus, measurements of the sediment nutrient concentrations before aquaculture introduction and continued monitoring thereafter would dramatically improve the certifiers’ abilities to assess the impact of nutrient loading from aquaculture on lake ecosystems.

Finally, addressing challenges associated with assessing the impact of net-pen aquaculture on inland waters has the potential to not only improve the sustainability of the aquaculture industry, but also provides an opportunity to improve data coverage on freshwater ecosystems (Fadum et al. *in review*). Adopting open data practices and making the pelagic data generated in the monitoring process publicly available would be particularly beneficial in regions of the globe that have scarce (due to existence or accessibility) ecological data. Such data is essential to understanding ecosystem change and regime shifts in globally distributed lakes (Gilarranz et al. 2022, Hernández Martínez de la Riva et al. 2023), a current, major challenge in assessing global freshwater sustainability.

## Conclusions

In this manuscript we compared the relative nutrient loading of a net-pen aquaculture operation to the principal watershed nutrient inputs for a large tropical lake. We found that the annual loading of both N (85.8%) and P (95.2%) from the aquaculture inputs dwarfed the annual nutrient inputs to Lake Yojoa from the watershed. The net-pen aquaculture practice in the center of Lake Yojoa had obtained multiple certifications that indicate the production of Tilapia was conducted sustainably. However, previous research in Lake Yojoa has shown a rapid (<40 year), nutrient driven, shift in its trophic state from a primarily oligo/mesotrophic ecosystem in the early 1980’s to a meso/eutrophic ecosystem with periodic hypereutrophic conditions in the early 2020s (Fadum and Hall 2022). This previously described shift in Lake Yojoa’s trophic state driven by reactive N combined with the analysis of relative nutrient inputs presented here strongly suggests that the aquaculture operation is the principal driver of the deterioration of Lake Yojoa’s ecosystem, despite the aquaculture operator’s compliance with top sustainability standards at the time of assessment. We conclude that the current best practices in sustainable net-pen aquaculture are insufficient and that certifying agencies do not require the appropriate assessments of ecosystem impact to ensure the sustainability of the practice. In a world of increasing human stress on freshwater ecosystems, it is essential that stakeholders are able to provide accurate indicators of best practices that consumers can rely on when making their purchasing decisions. Trusted certifiers must be able to provide ecologically relevant guidance to farm operators. This is currently not the case for Tilapia produced in net-pen farms. The failure to create such standards will result in the further deterioration of global freshwater resources and increasing stress on the ecosystem services on which local communities depend.

## Data availability statement

All code used for all data analysis and figure generation is available upon reasonable request. Nutrient loading data provided by Regal Springs Tilapia is available in Supplemental Text 4.

## Acknowledgments

This work was made possible by the collaborative efforts of Asociación de Municipios del Lago de Yojoa y su Área de Influencia (AMUPROLAGO) and supported in part by NSF RAPID (NSF DEB Award# 2120441, awarded to EKH). Additional financial support was provided by the Department of Ecosystem Science and Sustainability and the Graduate Degree Program in Ecology at Colorado State University. JMF was supported by the Simons Foundation (Award # 993455) during the final stages of manuscript preparation and submission.

## References

1. Amorim, L. F., Martins, J. R. S., Nogueira, F. F., Silva, F. P., Duarte, B. P. S., Magalhães, A. A. B., & Vinçon-Leite, B. (2021). Hydrodynamic and ecological 3D modeling in tropical lakes. SN Applied Sciences, 3(4), 444. 10.1007/s42452-021-04272-6

2. Boyd, C. E., McNevin, A. A., Clay, J., & Johnson, H. M. (2005). Certification Issues for Some Common Aquaculture Species. Reviews in Fisheries Science, 13(4), 231–279.

3. Bruce, L. C., Frassl, M. A., Arhonditsis, G. B., Gal, G., Hamilton, D. P., Hanson, P. C., Hetherington, A. L., Melack, J. M., Read, J. S., Rinke, K., Rigosi, A., Trolle, D., Winslow, L., Adrian, R., Ayala, A. I., Bocaniov, S. A., Boehrer, B., Boon, C., Brookes, J. D., … Hipsey, M. R. (2018). A multi-lake comparative analysis of the General Lake Model (GLM): Stress-testing across a global observatory network. Environmental Modelling & Software, 102, 274–291. 10.1016/j.envsoft.2017.11.016

4. Chow, V. T. (1959). Open-channel hydraulics. McGraw-Hill Book Co., New York, N.Y.

5. Coveney, M. F., Lowe, E. F., Battoe, L. E., Marzolf, E. R., & Conrow, R. (2005). Response of a eutrophic, shallow subtropical lake to reduced nutrient loading. Freshwater Biology, 50(10), 1718– 1730. 10.1111/j.1365-2427.2005.01435.x

6. Dentener, F.J. (2006). Global Maps of Atmospheric Nitrogen Deposition, 1860, 1993, and 2050. ORNL DAAC, Oak Ridge, Tennessee, USA. 10.3334/ORNLDAAC/830

7. Dingman, S.L. (2002) Physical Hydrology. 2nd Edition, Prentice Hall, Upper Saddle River, 646 p.

8. Dodds, W.K. (2006). Eutrophication and trophic state in rivers and streams. Limnology & Oceanography, 51(1), 671–680.

9. Dörnhöfer, K., & Oppelt, N. (2016). Remote sensing for lake research and monitoring – Recent advances. Ecological Indicators, 64, 105–122. 10.1016/j.ecolind.2015.12.009

10. Fadum, J. M., & Hall, E. K. (2022). The interaction of physical structure and nutrient loading drives ecosystem change in a large tropical lake over 40 years. Science of The Total Environment, 830, 154454. 10.1016/j.scitotenv.2022.154454

11. Fadum, J. M., & Hall, E. K. (2023). Nitrogen is unlikely to consistently limit primary productivity in most tropical lakes. Ecosphere, 14(3), e4451. 10.1002/ecs2.4451

12. Fadum, J. M., Borton, M. A., Daly, R. A., Wrighton, K. C., & Hall, E. K. (2023). Dominant nitrogen metabolisms of a warm, seasonally anoxic freshwater ecosystem revealed using genome resolved metatranscriptomics (p. 2023.08.22.554355). bioRxiv. 10.1101/2023.08.22.554355

13. Fadum, J. M., Waters, M. N., & Hall, E. K. (2023). Trophic state resilience to hurricane disturbance of Lake Yojoa, Honduras. Scientific Reports, 13(1), Article 1. 10.1038/s41598-023-32712-3

14. FAO (2022). *The State of World Fisheries and Aquaculture* 2022. FAO. 10.4060/cc0461en

15. Gilarranz, L. J., Narwani, A., Odermatt, D., Siber, R., & Dakos, V. (2022). Regime shifts, trends, and variability of lake productivity at a global scale. Proceedings of the National Academy of Sciences, 119(35), e2116413119. 10.1073/pnas.2116413119

16. Godsey, S. E., Kirchner, J. W., & Clow, D. W. (2009). Concentration–discharge relationships reflect chemostatic characteristics of US catchments. Hydrological Processes, 23(13), 1844–1864. 10.1002/hyp.7315

17. Gücker, B., Brauns, M., & Pusch, M. T. (2006). Effects of wastewater treatment plant discharge on ecosystem structure and function of lowland streams. Journal of the North American Benthological Society, 25(2), 313–329. 10.1899/0887-3593(2006)25[313:EOWTPD]2.0.CO;2

18. Hernández Martínez de la Riva, A., Harper, M., Rytwinski, T., Sahdra, A., Taylor, J. J., Bard, B., Bennett, J. R., Burton, D., Creed, I. F., Haniford, L. S. E., Hanna, D. E., Harmsen, E. J., Robichaud, C. D., Smol, J. P., Thapar, M., & Cooke, S. J. (2023). Tipping points in freshwater ecosystems: An evidence map. Frontiers in Freshwater Science, 1. https://www.frontiersin.org/articles/10.3389/ffwsc.2023.1264427

19. Imberger, J. (1998). Flux Paths in a Stratified Lake. In Physical Processes in Lakes and Oceans (pp. 1–17). American Geophysical Union (AGU). 10.1029/CE054p0001

20. Johnson, N. M., Likens, G. E., Bormann, F. H., Fisher, D. W., & Pierce, R. S. (1969). A Working Model for the Variation in Stream Water Chemistry at the Hubbard Brook Experimental Forest, New Hampshire. Water Resources Research, 5(6), 1353–1363. 10.1029/WR005i006p01353

21. Li, Y., Acharya, K., Stone, M. C., Yu, Z., Young, M. H., Shafer, D. S., Zhu, J., Gray, K., Stone, A., Fan, L., Tang, C., & Warwick, J. (2011). Spatiotemporal patterns in nutrient loads, nutrient concentrations, and algal biomass in Lake Taihu, China. Lake and Reservoir Management, 27(4), 298–309. 10.1080/07438141.2011.610560

22. Luo, L., Zhang, H., Luo, C., McBridge, C., Muraoka, K., Zhou, H., Hou, C., Liu, F., & Li, H. (2022). Tributary Loadings and Their Impacts on Water Quality of Lake Xingyun, a Plateau Lake in Southwest China. Water, 14(8), Article 8. 10.3390/w14081281

23. McClain, M. E., Boyer, E. W., Dent, C. L., Gergel, S. E., Grimm, N. B., Groffman, P. M., Hart, S. C., Harvey, J. W., Johnston, C. A., Mayorga, E., McDowell, W. H., & Pinay, G. (2003). Biogeochemical Hot Spots and Hot Moments at the Interface of Terrestrial and Aquatic Ecosystems. Ecosystems, 6(4), 301–312.

24. Meals, D. W., & Dressing, S.A. (2008). Surface Water Flow Measurement for Water Quality Monitoring Projects. Tech Notes #9, September 2013. Prepared for U.S. Environmental Protection Agency, by Tetra Tech, Inc., Fairfax, VA. Accessed March 16, 2016. https://www.epa.gov/pollutedrunoff-nonpoint-source-pollution/nonpoint-source-monitoring-technical-notes.

25. Mooney, R. J., Stanley, E. H., Rosenthal, W. C., Esselman, P. C., Kendall, A. D., & McIntyre, P. B. (2020). Outsized nutrient contributions from small tributaries to a Great Lake. Proceedings of the National Academy of Sciences, 117(45), 28175–28182. 10.1073/pnas.2001376117

26. Naylor, R. L., Hardy, R. W., Buschmann, A. H., Bush, S. R., Cao, L., Klinger, D. H., Little, D. C., Lubchenco, J., Shumway, S. E., & Troell, M. (2021). A 20-year retrospective review of global aquaculture. Nature, 591(7851), Article 7851. 10.1038/s41586-021-03308-6

27. North, R. L., Winter, J. G., & Dillon, P. J. (2013). Nutrient indicators of agricultural impacts in the tributaries of a large lake. Inland Waters, 3(2), 221–234. 10.5268/IW-3.2.565

28. Preston, S. D., Alexander, R. B., Schwarz, G. E., & Crawford, C. G. (2011). Factors Affecting Stream Nutrient Loads: A Synthesis of Regional SPARROW Model Results for the Continental United States1. JAWRA Journal of the American Water Resources Association, 47(5), 891–915. 10.1111/j.1752-1688.2011.00577.x

29. Rector, M. E., Filgueira, R., Bailey, M., Walker, T. R., & Grant, J. (2023). Sustainability outcomes of aquaculture eco-certification: Challenges and opportunities. Reviews in Aquaculture, 15(2), 840– 852. 10.1111/raq.12763

30. Reinaldo Finkler, N., Tromboni, F., Boëchat, I. G., Gücker, B., & Gasparini Fernandes Cunha, D. (2018). Nitrogen and Phosphorus Uptake Dynamics in Tropical Cerrado Woodland Streams. Water, 10(8), Article 8. 10.3390/w10081080

31. Revelle, W. (2024). psych: Procedures for Psychological, Psychometric, and Personality Research. Northwestern University, Evanston, Illinois. R package version 2.4.1, https://CRAN.R-project.org/package=psych.

32. Riveros-Iregui, D. A., Covino, T. P., & González-Pinzón, R. (2018). The importance of and need for rapid hydrologic assessments in Latin America. Hydrological Processes, 32(15), 2441–2451. 10.1002/hyp.13163

33. Robertson, D. M., & Saad, D. A. (2011). Nutrient Inputs to the Laurentian Great Lakes by Source and Watershed Estimated Using SPARROW Watershed Models1. JAWRA Journal of the American Water Resources Association, 47(5), 1011–1033. 10.1111/j.1752-1688.2011.00574.x

34. Sterner, R. W., & Elser, J. J. (2017). Ecological Stoichiometry: The Biology of Elements from Molecules to the Biosphere. In Ecological Stoichiometry. Princeton University Press. 10.1515/9781400885695

35. Tromboni, F., & Dodds, W. K. (2017). Relationships Between Land Use and Stream Nutrient Concentrations in a Highly Urbanized Tropical Region of Brazil: Thresholds and Riparian Zones. Environmental Management, 60(1), 30–40. 10.1007/s00267-017-0858-8

36. Turner, R. E., Rabalais, N. N., Justic’, D., & Dortch, Q. (2003). Global patterns of dissolved N, P and Si in large rivers. Biogeochemistry, 64(3), 297–317. 10.1023/A:1024960007569

37. Wickham, H. (2009). GGPLOT2: Elegant graphics for data analysis. Springer.

38. Wymore, A. S., Brereton, R. L., Ibarra, D. E., Maher, K., & McDowell, W. H. (2017). Critical zone structure controls concentration-discharge relationships and solute generation in forested tropical montane watersheds. Water Resources Research, 53(7), 6279–6295. 10.1002/2016WR020016

