## Supplemental Text 1 for "Nutrient loading from a sustainably certified aquaculture operation dwarfs annual nutrient inputs from a large multi-use watershed, Lake Yojoa, Honduras"

### Supplemental Text 1: Hydrology characterization

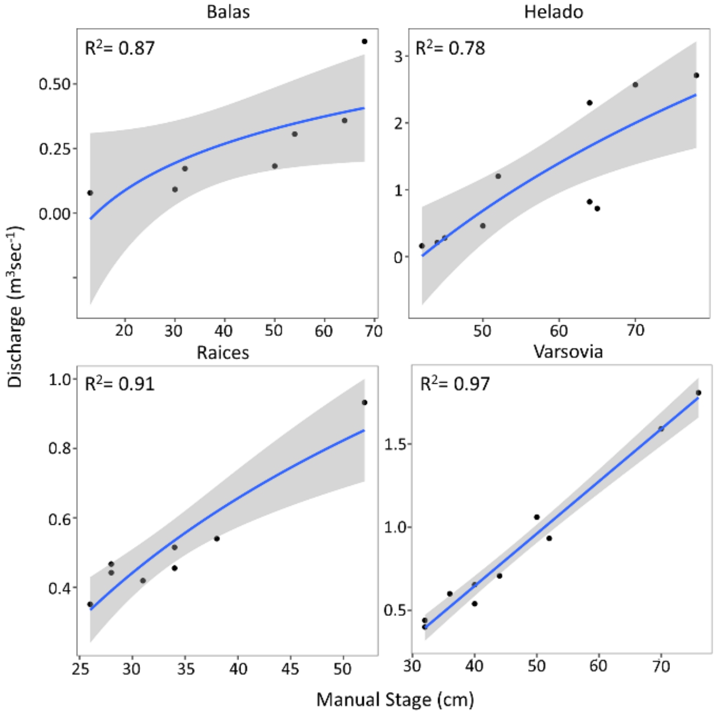


**Figure S1.** Rating curves with coefficient of determination with linear regression (Varsovia) and logistic regression (Balas, Helado and Raices).

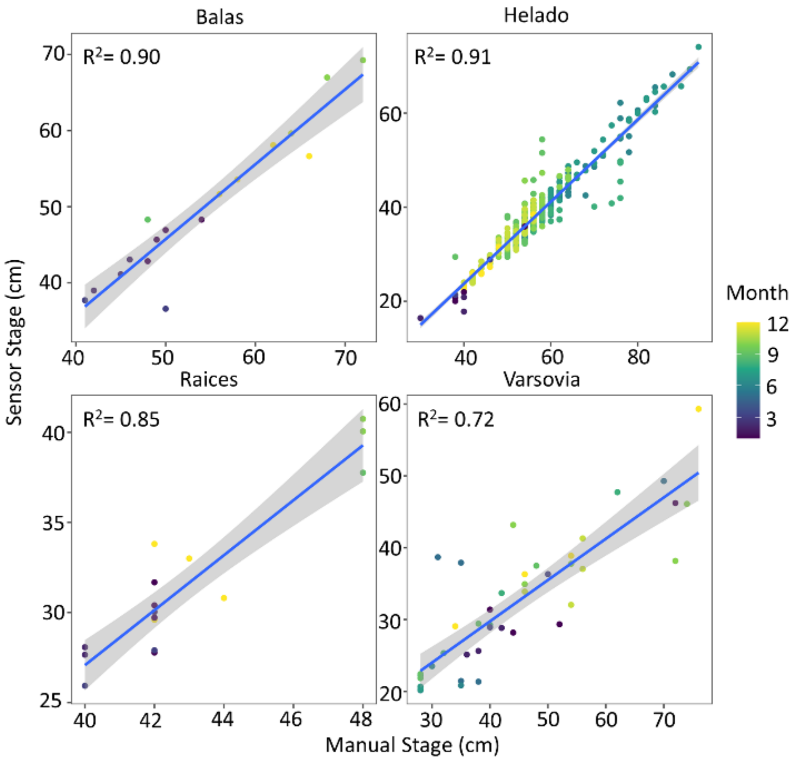


**Figure S2.** Manual stage observations vs. calculated sensor stage at four of the major tributaries with coefficient of determination with linear regression.
