## Supplemental Text 2 for "Nutrient loading from a sustainably certified aquaculture operation dwarfs annual nutrient inputs from a large multi-use watershed, Lake Yojoa, Honduras"

### Supplemental Text 2: Sampling Frequently Analysis

For the tributaries where we were able to install pressure transducers, we modeled discharge at 15 minute intervals. We then tested how frequently discharged would be measured to identify a mean discharge value that was with 10 percent of the true (i.e., pressure transducer modeled value). We tested the following scenarios:  Apply the observation taken: 1) on the first day of each month to the whole month (i.e. sampling frequency = 12), 2) twice a month (i.e. sampling frequency = 24), 3) weekly (i.e. sampling frequency = 52), 4) daily (i.e. sampling frequency = 365), 5) twice daily at 8:00 and 16:00 (i.e. sampling frequency = 730), 6) three times daily at 6:00, 12:00, and 18:00 (i.e. sampling frequency = 1095), 7) four times daily at 5:00, 10:00, 15:00 and 20:00 (i.e. sampling frequency = 1460), and 8) hourly (i.e. sampling frequency = 8760).

Our analysis suggests that less frequency discharge analysis yields comparable results for all tributaries. Twice daily measurements would have been sufficient for accurately calculating discharge at most tributaries during most months.

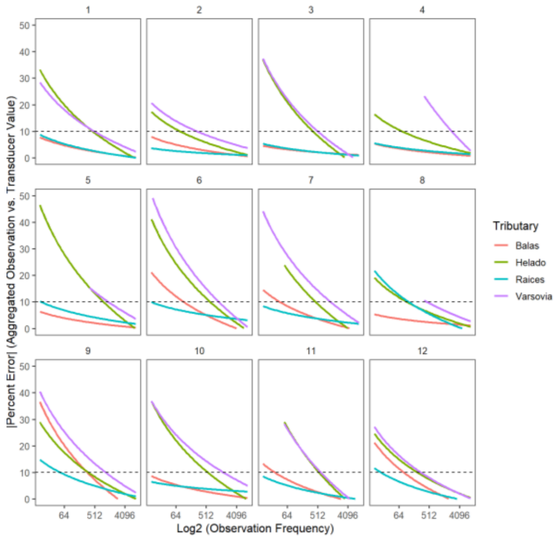


**Figure S3.** Sampling frequency vs. absolute error from 'true' discharge.

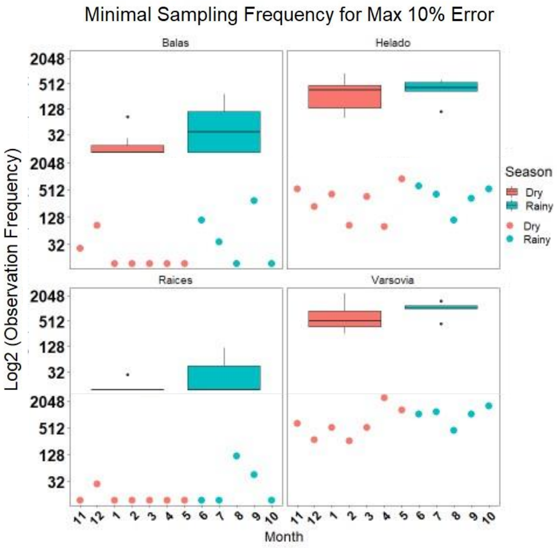


**Figure S4.** Sampling frequency required to achieve <10% error in each month, divided into rainy and dry seasons.
