## Supplemental Text 3 for "Nutrient loading from a sustainably certified aquaculture operation dwarfs annual nutrient inputs from a large multi-use watershed, Lake Yojoa, Honduras"

**Supplemental Text 3: Predictions of aquaculture nutrient loading after 2013**

Using Google Earth, we quantified the number of net-pens operated by Regal Springs Tilapia in Lake Yojoa since 2013 (i.e., spanning the years we are unable to obtain loading estimates from operator data). We assume a linear relationship between nutrient loading and use a per pen loading estimate (Table 2).

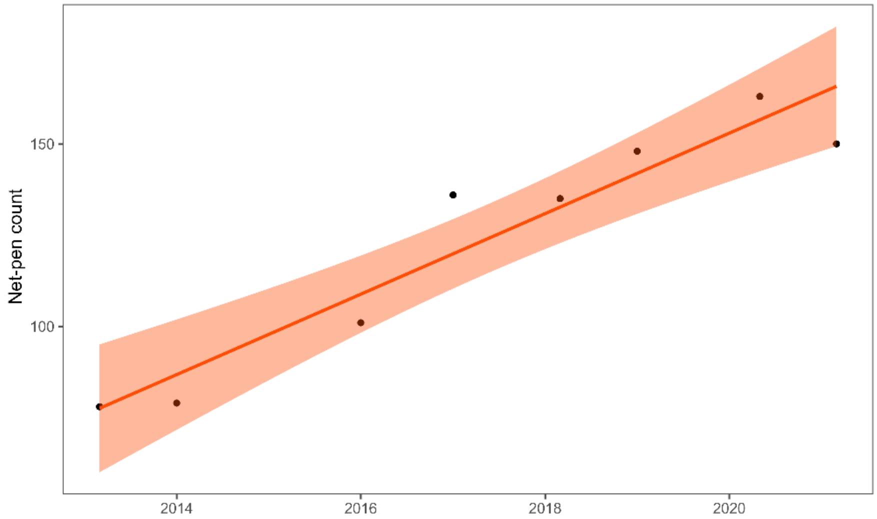


**Figure S5**. Number of Regal Springs Tilapia pens in Lake Yojoa 2013-2021.

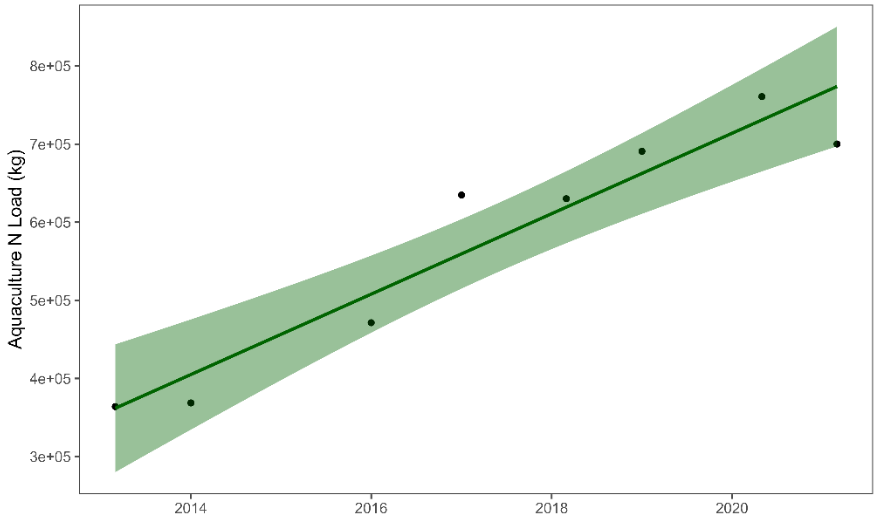


**Figure S6.** Estimated N loading based on net-pens operated for 2013-2021.

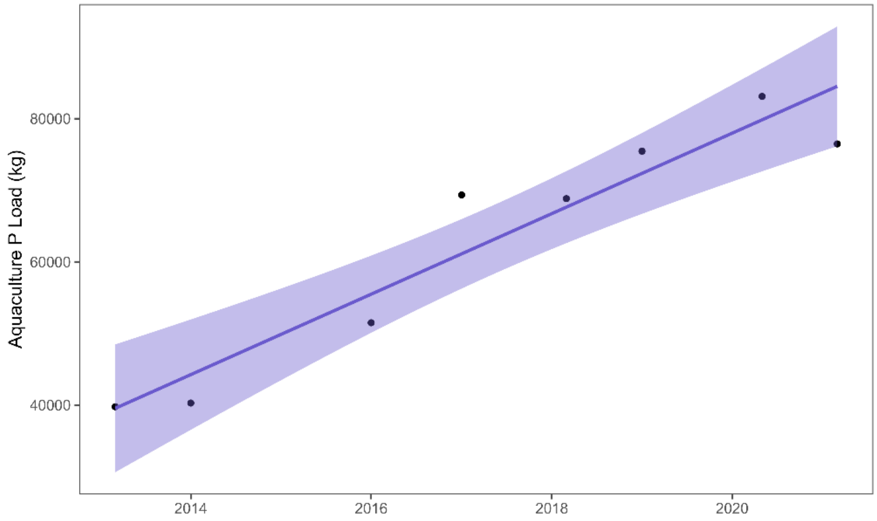


**Figure S7.** Estimated P loading based on net-pens operated for 2013-2021.
