## Supplemental Text 4 for "Nutrient loading from a sustainably certified aquaculture operation dwarfs annual nutrient inputs from a large multi-use watershed, Lake Yojoa, Honduras"

### Supplemental Text 4: Calculating Nutrient Load from Regal Spring Tilapia

Regal Spring Tilapia provided information on the following: 1) the amount of various feeds supplied to fish pens in Lake Yojoa, 2) the N and P content of those feed formulas, 3) the mass of harvested biomass and 4) the N and P content of harvested biomass.

**Table S1.** N and P content (in %) of various feed formulas and quantity of feed proved per month (in kg).

| | **Aquaxcel 45%** | **Alcon 40%** | **Alcon 35%** | **Alcon 36%** | **Alcon 32%** | **Gisis-Expalsa 32%** | **Gisis-Expalsa 36%** | **Gisis-Expalsa 40%** | **Aquafeed 32 %** | **Aquafeed 36 %** | **Aquafeed 40 %** |
|---|---|---|---|---|---|---|---|---|---|---|---|
| N content | 7.2% | 6.40% | 5.60% | 5.80% | 5.20% | 5.12% | 5.76% | 6.41% | 5.12% | 5.76% | 6.41% |
| P content | 1.1% | 1.03% | 0.99% | 1.00% | 1.01% | 0.84% | 0.90% | 0.87% | 0.84% | 0.90% | 87.00% |
| January | 29,396.91 | 20,322.19 | 64,139.04 | - | 141,932.35 | 158,181.21 | 55,180.89 | 16,908.85 | - | - | - |
| February | 14,570.99 | 17,236.80 | 5,851.44 | 19,278.00 | 44,362.08 | 273,498.12 | 55,408.60 | 43,930.71 | - | - | - |
| March | 7,734.33 | 2,812.32 | 13,653.36 | 9,072.45 | 72,213.57 | 295,409.27 | 116,317.56 | 14,067.95 | - | - | - |
| April | 22,080.79 | 1,950.48 | 9,298.80 | - | 133,630.56 | 353,818.43 | 67,363.23 | 61,044.13 | - | - | - |
| May | 10,878.24 | 99,111.60 | 125,193.60 | - | 398,623.68 | 132,454.38 | - | 3,228.72 | - | - | - |
| June | 7,734.33 | 61,598.88 | 183,435.84 | - | 585,234.27 | 141,409.35 | 2,272.99 | - | - | - | - |
| July | 22,055.85 | 35,290.08 | 159,032.16 | 7,076.16 | 752,885.28 | 178,181.79 | - | - | - | - | - |
| August | 9,605.89 | 46,267.65 | 75,614.67 | - | 420,804.72 | 104,545.27 | - | - | 269,846.64 | - | - |
| September | 16,167.66 | 9,117.36 | 1,360.80 | - | - | 51,445.04 | 107,818.00 | 71,138.54 | 586,005.84 | - | - |
| October | 1,197.50 | - | 38,556.00 | - | - | 15,646.02 | 19,163.47 | 13,405.69 | 588,409.92 | 116,575.20 | 40,824.00 |
| November | - | - | - | - | - | - | 6,086.40 | 1.36 | 461,401.92 | 118,026.72 | 21,364.56 |
| December | - | - | - | - | - | - | 6,899.71 | - | 707,661.36 | 87,318.00 | 24,449.04 |

**Table S2.** N additions calculated by mass of feed supplied multiplied by N content by percentage. All values are provided in kg.

| | Aquaxcel 45% | Alcon 40% | Alcon 35% | Alcon 36% | Alcon 32% | Gisis-Expalsa 32% | Gisis-Expalsa 36% | Gisis-Expalsa 40% | Aquafeed 32 % | Aquafeed 36 % | Aquafeed 40 % |
|---|---|---|---|---|---|---|---|---|---|---|---|
| January | 2,116.58 | 1,300.62 | 3,591.79 | - | 7,380.48 | 8,098.88 | 3,178.42 | 1,083.86 | - | - | - |
| February | 1,049.11 | 1,103.16 | 327.68 | 1,118.12 | 2,306.83 | 14,003.10 | 3,191.54 | 2,815.96 | - | - | - |
| March | 556.87 | 179.99 | 764.59 | 526.20 | 3,755.11 | 15,124.95 | 6,699.89 | 901.76 | - | - | - |
| April | 1,589.82 | 124.83 | 520.73 | - | 6,948.79 | 18,115.50 | 3,880.12 | 3,912.93 | - | - | - |
| May | 783.23 | 6,343.14 | 7,010.84 | - | 20,728.43 | 6,781.66 | - | 206.96 | - | - | - |
| June | 556.87 | 3,942.33 | 10,272.41 | - | 30,432.18 | 7,240.16 | 130.92 | - | - | - | - |
| July | 1,588.02 | 2,258.57 | 8,905.80 | 410.42 | 39,150.03 | 9,122.91 | - | - | - | - | - |
| August | 691.62 | 2,961.13 | 4,234.42 | - | 21,881.85 | 5,352.72 | - | - | 13,816.15 | - | - |
| September | 1,164.07 | 583.51 | 76.20 | - | - | 2,633.99 | 6,210.32 | 4,559.98 | 30,003.50 | - | - |
| October | 86.22 | - | 2,159.14 | - | - | 801.08 | 1,103.82 | 859.31 | 30,126.59 | 6,714.73 | 2,616.82 |
| November | - | - | - | - | - | - | 350.58 | 0.09 | 23,623.78 | 6,798.34 | 1,369.47 |
| December | - | - | - | - | - | - | 397.42 | - | 36,232.26 | 5,029.52 | 1,567.18 |

**Table S3.** P additions calculated by mass of feed supplied multiplied by N content by percentage. All values are provided in kg.

| | Aquaxcel 45% | Alcon 40% | Alcon 35% | Alcon 36% | Alcon 32% | Gisis-Expalsa 32% | Gisis-Expalsa 36% | Gisis-Expalsa 40% | Aquafeed 32 % | Aquafeed 36 % | Aquafeed 40 % |
|---|---|---|---|---|---|---|---|---|---|---|---|
| January | 323.37 | 209.32 | 634.98 | - | 1,433.52 | 1,328.72 | 496.63 | 147.11 | - | - | - |
| February | 160.28 | 177.54 | 57.93 | 192.78 | 448.06 | 2,297.38 | 498.68 | 382.20 | - | - | - |
| March | 85.08 | 28.97 | 135.17 | 90.72 | 729.36 | 2,481.44 | 1,046.86 | 122.39 | - | - | - |
| April | 242.89 | 20.09 | 92.06 | - | 1,349.67 | 2,972.07 | 606.27 | 531.08 | - | - | - |
| May | 119.66 | 1,020.85 | 1,239.42 | - | 4,026.10 | 1,112.62 | - | 28.09 | - | - | - |
| June | 85.08 | 634.47 | 1,816.01 | - | 5,910.87 | 1,187.84 | 20.46 | - | - | - | - |
| July | 242.61 | 363.49 | 1,574.42 | 70.76 | 7,604.14 | 1,496.73 | - | - | - | - | - |
| August | 105.66 | 476.56 | 748.59 | - | 4,250.13 | 878.18 | - | - | 2,266.71 | - | - |
| September | 177.84 | 93.91 | 13.47 | - | - | 432.14 | 970.36 | 618.91 | 4,922.45 | - | - |
| October | 13.17 | - | 381.70 | - | - | 131.43 | 172.47 | 116.63 | 4,942.64 | 1,049.18 | 355.17 |
| November | - | - | - | - | - | - | 54.78 | 0.01 | 3,875.78 | 1,062.24 | 185.87 |
| December | - | - | - | - | - | - | 62.10 | - | 5,944.36 | 785.86 | 212.71 |

**Table S4.** N and P loading calculated by cumulative N and P additions (Tables S2 and S3, respectively), less N and P in biomass in harvested Tilapia. Biomass is assumed to be 2.25% N by weight and 0.80% P by weight. All values are provided in kg.

| | **Biomass harvested** | **Biomass N** | **N in feed supplies** | **N loading** | **Biomass P** | **P in feed supplied** | **P loading** |
|---|---|---|---|---|---|---|---|
| January | 490,337.39 | 11,032.59 | 26,750.62 | 15,718.03 | 3,922.70 | 4,573.63 | 650.94 |
| February | 219,173.32 | 4,931.40 | 25,915.50 | 20,984.10 | 1,753.39 | 4,214.84 | 2,461.46 |
| March | 300,827.90 | 6,768.63 | 28,509.36 | 21,740.73 | 2,406.62 | 4,719.98 | 2,313.36 |
| April | 466,987.96 | 10,507.23 | 35,092.72 | 24,585.50 | 3,735.90 | 5,814.13 | 2,078.23 |
| May | 226,337.98 | 5,092.60 | 41,854.27 | 36,761.67 | 1,810.70 | 7,546.73 | 5,736.03 |
| June | 372,947.98 | 8,391.33 | 52,574.87 | 44,183.54 | 2,983.58 | 9,654.72 | 6,671.14 |
| July | 532,173.41 | 11,973.90 | 61,435.75 | 49,461.84 | 4,257.39 | 11,352.15 | 7,094.76 |
| August | 595,681.62 | 13,402.84 | 48,937.89 | 35,535.05 | 4,765.45 | 8,725.83 | 3,960.37 |
| September | 1,080,198.86 | 24,304.47 | 45,231.57 | 20,927.10 | 8,641.59 | 7,229.08 | (1,412.51) |
| October | 388,061.40 | 8,731.38 | 44,467.69 | 35,736.31 | 3,104.49 | 7,162.39 | 4,057.90 |
| November | 461,157.19 | 10,376.04 | 32,142.25 | 21,766.21 | 3,689.26 | 5,178.68 | 1,489.42 |
| December | 292,181.29 | 6,574.08 | 43,226.39 | 36,652.31 | 2,337.45 | 7,005.02 | 4,667.57 |
| | | | **TOTAL:** | **364,052.38** | | **TOTAL:** | **39,768.67** |
