## Supplemental Text 5 for "Nutrient loading from a sustainably certified aquaculture operation dwarfs annual nutrient inputs from a large multi-use watershed, Lake Yojoa, Honduras"

**Supplemental Text 5: Comparison of dissolved and particulate fraction of tributary nitrogen**

In June 2019 water from Lake Yojoa's major tributaries was filtered through Whatman glass microfiber filters (Grade GF/F, pore size = 0.7 μm). Total dissolved nitrogen (TDN) in filtered water was measured using a Shimadzu TOC-L (Shimadzu Scientific Instruments, Inc.). Total nitrogen (TN) on filters was measured on a Delta V IRMS coupled to a Costech ECS 4010 elemental analyzer. All analyses were performed at EcoCore at Colorado State University (Fort Collins, CO, USA).

We found that the dissolved N fraction in Lake Yojoa's tributaries exceeds the particulate (> 0.7 μm) fraction. We also assessed the $\delta^{15}N$ of the particulate fraction and found that Raices was enriched relative to other tributaries, likely reflective of agricultural and municipal waste contributions.

**Table S5.** Dissolved and particulate fractions of nitrogen in Lake Yojoa's six main tributaries and $\delta^{15}N$ of the particulate pool.

| | TN ($\mu gL^{-1}$) | TN- $\delta^{15}N$ (‰) | TDN ($\mu gL^{-1}$) |
|---|---|---|---|
| Balas | 6.45 | 19.14 | 1056.13 |
| Cacao | 10.00 | 10.33 | 367.88 |
| Helado | 176.00 | 12.26 | 588.50 |
| Raices | 140.00 | 29.43 | 1815.75 |
| Varsovia | 15.00 | 2.24 | 219.25 |
| Yure | 60.00 | 4.75 | 178.63 |
